# Metabolomics unveiled the impact of alterations in nucleotide glucose metabolism on palmitic acid-induced TLR4 activation in THP-1 macrophages

**DOI:** 10.1101/2024.01.02.573974

**Authors:** Niting Wu, Xiaodong Ran, Jiawei Wang, Pengbo Wang, Lin Li, Yuanting She, Yadan Luo, Xiaohui Li, Yi Jia, Yan Huang

**Author notes:** Correspondence: Shapingba, Chongqing, China; (Y.H.); (Y.J.); (HX.L.). (Y.H., Y.J. and X.L.); (Y.H., Y.J. and X.L.). These authors contributed equally to this work (Niting Wu, Xiaodong Ran).

## Abstract

**OBJECTIVE:** The association of palmitic acid with macrophage inflammation and its promotion of the expression of inflammatory factors through TLR4 have been demonstrated. It has been observed that immature TLR4 localizes to the endoplasmic reticulum and Golgi apparatus, necessitating glycosylation for migration to the cell membrane. The objective of this study was to identify potential biomarkers associated with N-glycosylation subsequent to the activation of TLR4 inflammatory signaling in human macrophages by palmitic acid.

**APPROACH AND RESULTS:** The co-cultivation of palmitic acid with THP-1 macrophages was conducted for a duration of 24 hours, followed by the collection of cell extracts for subsequent metabolomic and lipidomic analyses using high performance liquid chromatography-tandem mass spectrometry. Multivariate and univariate statistical analyses were conducted to identify potential biomarkers, in accordance with established scientific protocols. The impact of palmitic acid on the TLR4 signaling pathway and macrophage N-glycosylation was assessed at various time points using Western blot analysis, immunofluorescence staining, Elisa assays, and chemical labeling techniques. The TLR4 inflammatory signaling pathway was examined in macrophages at different time points, revealing that PA induced the upregulation of MyD88 and TRAF6 expression as well as NF-κB phosphorylation, indicating the activation of classical NF-κB signaling. After 24 hours, TLR4 translocated from the cell membrane to the cytoplasm and initiated internalization, accompanied by significant colocalization with GalAz in the cytosol. In addition, the metabolites in the cell extract were found to be altered in both the control and model groups. Significantly alterations in two N-glycosylation related metabolites were observed in the model group, including guanosine diphosphate-L-fucose and uridine diphosphate-N-acetylglucosamine/uridine diphosphate-N-acetylgalactosamine.

**CONCLUSIONS:** THP-1 macrophages incubated with palmitic acid exhibited a distinct metabolomic profile compared to the control group. Our findings suggest that metabolomics analysis holds promise in identifying disease-specific biomarkers for diagnosing fatty acid-induced inflammatory responses in macrophages.

## INTRODUCTION

Obesity has emerged as a chronic ailment, exhibiting an alarming surge in global prevalence in recent years^1,2^. According to the World Health Organization, approximately 2 billion adults are affected by overweight, while 650 million individuals suffer from obesity. If current patterns persist, it is projected that by 2025, the number of overweight adults will escalate to 2.7 billion and surpassing one billion for obesity^3,4^. Clinical evidence suggests that the prevalence of obesity is on the rise, irrespective of race, socioeconomic status, or age^5^. Plasma concentrations of Free fatty acids (FFA) are found to be elevated in individuals who are overweight or obese ^6–8^. The excessive intake of saturated fatty acids and the presence of metabolic disorders in obese patients contribute to an elevation in blood fatty acid levels. This leads to an inflammatory response, which is an important factor in the development and progression of obesity-related diseases^9,10^. The FFA molecules constitute a highly intricate group, thus their impact on cellular functions is likely to exhibit significant variability, potentially inducing either pro-inflammatory or anti-inflammatory responses^11,12^. Palmitic acid (PA) is a kind of saturated fatty acid^12^ in human body with the molecular formula of C_16_H_32_O_2_. It constitutes 65% of the total saturated fatty acids present in the human body and accounts for approximately 28-32% of the overall fatty acid composition in serum^13,14^. The association between PA and inflammation has been extensively demonstrated in numerous studies^15,16^, with the promotion of inflammatory factor expression mediated through Toll-like receptor 4 (TLR4) ^17–19^. The role of macrophages in inflammation, repair, and regeneration is well-established^20^. The present study aimed to investigate the modulation of macrophage inflammatory response through PA stimulation. The question of whether PA functions as a direct TLR4 agonist or indirectly activates TLR4 remains open for discussion^21–26^. However, it is evident that TLR4 plays a pivotal role in the induction of macrophage inflammation by PA^27–30^. TLR4 is one of the ten Toll-like receptors identified in mammals that play a crucial role in innate immunity. In its immature state, TLR4 localizes to the endoplasmic reticulum (ER) and Golgi apparatus, requiring glycosylation for maturation and subsequent migration to the cell membrane^31,32^. The extracellular domain of Toll-like receptors is extensively glycosylated, featuring multiple N-linked glycosylation sites on the concave surface (FIG. 1). The N-linked glycosylation modification of human MD-2 is essential for the activation of nuclear factor-κB (NF-κB)mediated by human TLR4 ^33,34^.

**Figure 1.**
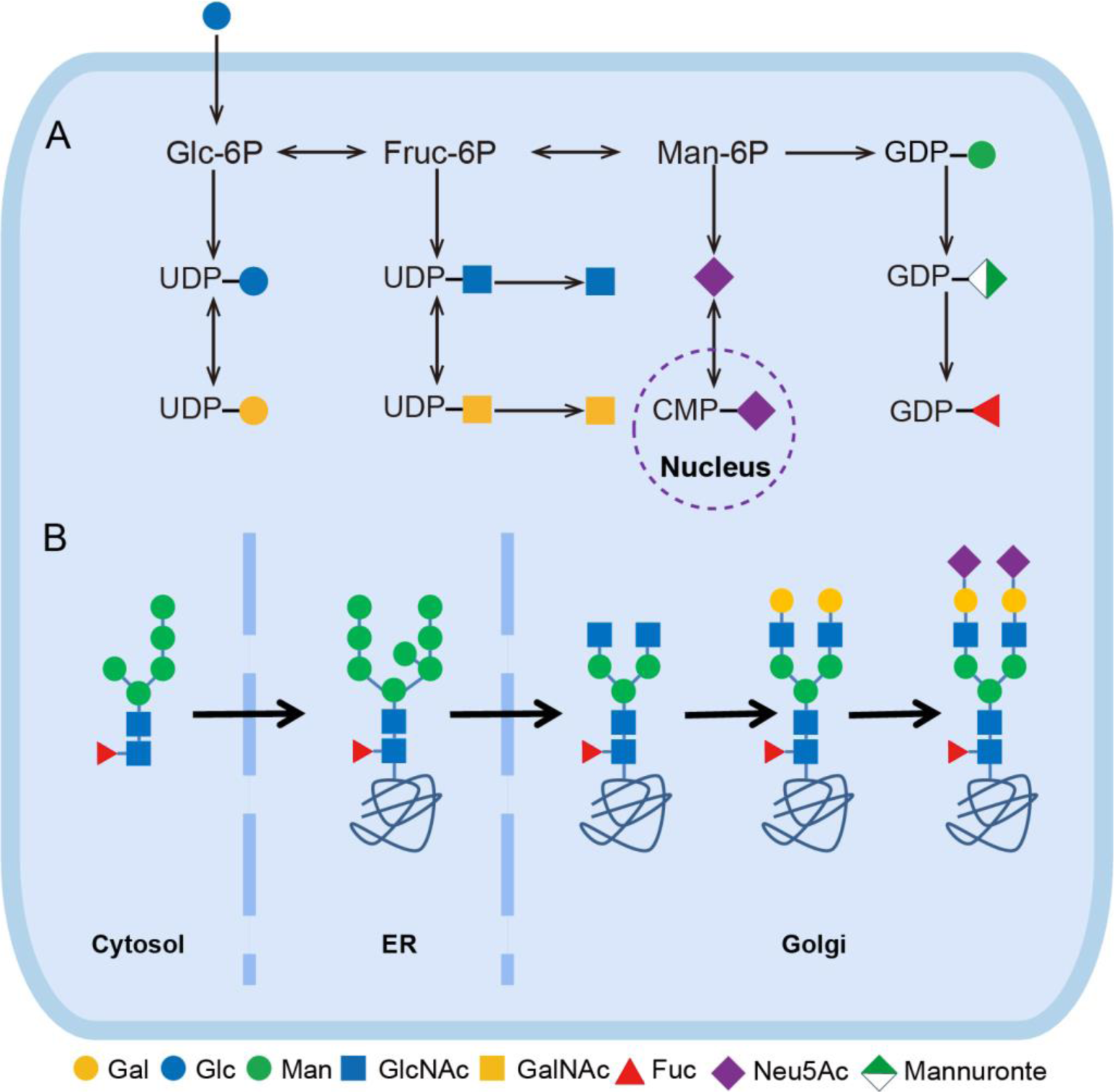
Nucleotide sugar synthesis and N-glycosylation in cells. (A) The pathway of intracellular biosynthesis of nucleotide sugars. (B) The cellular process of protein N-glycosylation.

Glycosylation is the most prevalent posttranslational modification of proteins. In general, glycosylation occurs in two primary forms, namely N-linked and O-linked, which are determined by the functional group of the amino acids to which the glycans are attached. The N-glycans are covalently attached to asparagine residues via amide linkages^35^. Sequence analysis indicates that human myeloid differentiation protein-2 (MD-2) possesses two potential N-linked glycosylation sites, while the ectodomain of human TLR4 contains nine^36^. It was found that MD-2 undergoes N-glycosylation, which is crucial for TLR4-mediated NF-κB activation. The process of N-linked glycosylation initiates with the attachment of a choline diphosphate-linked 7-sugar oligosaccharide to the ER. This oligosaccharide flips into the phosphorous endoplasmic reticulum ER into 14 sugar N-glycosylated oligosaccharide precursors. Subsequently, this lipid-linked oligosaccharide is separated from its phosphate polyphenol anchor and synchronized with the asparagine amino acid binding of the protein, with the removal of the three terminal glucose (Glc) moieties and the most central terminal mannose (Man) (Figure 1A)^37^. The process of glycosylation continues within the Golgi apparatus, where N-glycosylated glycoproteins undergo processing. Initially, in the cis-Golgi, Man residues are removed from these glycoproteins by a range of α-mannosidases to generate the Man5GlcNAc2Asn intermediate. This intermediate serves as a substrate for alpha-1,3-mannosyl-glycoprotein 2-beta-N-acetylglucosaminyl transferase (MGAT1) in the medial Golgi. The N-glycan loses the terminal two of five Man residues and acquires a second N-acetylglucosamine (GlcNAc), resulting in the formation of a complex N-glycan with two antennae. This structure can further undergo up to six branching events, with each branch being elongated by the addition of various sugars, including Galactose (Gal), GlcNAc, N-acetylgalactosamine (GalNAc), Fucose (Fuc), N-Acetylneuraminic acid (Neu5Ac, also known as sialic acid), as well as disaccharide units (Figure 1B) ^38^.Glycosylation is regulated by a variety of factors, including glycosyltransferases, gene expression, localization, substrate availability, acceptor and donor substrates such as nucleotide sugars. The substrates for glycosylation typically consist of monosaccharides that are linked to specific nucleoside-phosphates, including uridine diphosphate galactose (UDP-Gal), uridine diphosphate-N-acetylglucosamine (UDP-GlcNAc), guanosine diphosphate-L-fucose (GDP-L-Fuc), cytidine monophosphate-N-acetylneuraminic acid (CMP-Neu5Ac), etc. Nucleotide-sugars are generally produced in the cytoplasm, except for CMP-Neu5A, which is biosynthesized within the nucleus. The role of N-glycosylation in macrophage-associated inflammatory responses has been extensively investigated^39–41^, but the role of cellular nucleotide sugar content in inflammation remains limited. The regulation of nucleotide sugars as N-glycosylation substrates in the activation-induced dimerization of human macrophage TLR4 is a question of great interest for the inflammatory response of macrophages stimulated by PA. However, no relevant research results have been presented.

We conducted metabolomic analysis of human macrophages treated with PA, aiming to gain a deeper understanding of the role played by nucleotide sugars in pathological changes that occur following PA treatment, specifically their association with TLR4-activated N-glycosylation. We also used molecular analyses to elucidate the effects of PA treatment on TLR4 signaling in human macrophages following TLR4 activation. We hereby provide evidence of a nucleotide sugar response in human macrophages induced by PA. Nucleotide sugars have been identified as key regulators of TLR4 signaling in PA-activated macrophages. Therefore, intracellular nucleotide sugars can be considered a significant metabolic indicator in the PA-induced inflammatory response within human macrophages.

## METHODS

### Cell culture

A human monocyte leukemia cell line (THP-1) was obtained from ATCC, The cells were cultured in RPMI 1640 medium supplemented with 10% FBS, 1% penicillin/streptomycin solution and 0.05mM β-mercaptoethanol (referred to hereafter as media) in a humidified incubator (37 °C/5% CO_2_). THP-1 cells were differentiated by incubating the cells with 100 ng/mL phorbol-12-myristate-13-acetate (PMA, HY-18739, Med Cham Express, USA) for 24 h, with the media changed on the next day followed by further incubation for 24 h. Palmitic acid (PA, 932787, J&K, China) was dissolved in dimethyl sulfoxide (DMSO). Cells were treated with 100 μM PA+0.4% DMSO for 0h, 12h, 24h, 36h and 48h.

### Metabolic Labeling of Cell and Immunofluorescence assays

THP-1 cells were cultured in RPMI 1640 medium with low glucose or containing 50 μM 2-azide-2-deoxy-D-galactose (GalAz, A886798, MACKLIN). Cells were treated with PA for 12 h, 24 h, 36 h, and 48 h and harvested according to the set time points. After washing with PBS, cells were incubated with PBS containing 0.5% FBS, 50 μM Fluor 488-alkyne (761621, Sigma, USA), 2.5 mM sodium ascorbate (A103539, Aladdin, China) and BTTAA-CuSO4 complex (50μM CuSO4, BTTAA/CuSO4 in a 6:1 molar ratio) at room temperature for 5min Then cells were fixed with 4% paraformaldehyde for 15 min^42,43^. After blocking and permeabilized with 0.1% Triton X-100, the samples were incubated with an antibody against TLR4 (66350, proteintech, China) overnight at 4 °C. Followed by incubation with the appropriate secondary antibody for 1h at room temperature. Finally, Nuclei were stained with DAPI and cell membranes were stained with FITC-conjugated wheat germ agglutinin (WGA, FL-1021, VectorLabs, USA) when necessary. The coverslips were mounted onto microscope slides and imaged under a confocal laser scanning microscope (Zeiss LSM 900).

### Flow Cytometry

The cells in each group were collected into tubes and washed in ice-cold phosphate-buffered saline (PBS) twice. Then stained with APC-conjugated TLR4 antibody (312816, BioLegend, USA) for 30 minutes at room temperature. The stained cells were analyzed by flow cytometry (BD-FACSVerse) and software (FlowJo). For each sample, the count in 10,000 cells was measured.

## ELISA

The Human TNF-α ELISA Kit (1117202, Dakewe, China) and Human IL-1β ELISA Kit (1110123, Dakewe, USA) were used to detected the expression of TNF-α and IL-1β in cell supernatant of each group. Each experiment was repeated at least three times.

### Western blot

We lysed the cells using a protein extraction reagent (C2978, Sigma, Germany) in the presence of protease and phosphatase inhibitor cocktail tablets (87785, Thermo, USA). Protein concentration was measured with a BCA Protein Assay Kit (P0010, Beyotime, China). Equal amounts of protein extracts were separated using 10% sodium dodecyl sulphate polyacrylamide gel electrophoresis. After blocking by 5% (w/v) skim milk (blocking buffer) for 1 h at room temperature, the PVDF (IPVH00010, Millipore, USA) membranes with protein on it were incubated with primary antibodies against TLR4 (38519, CST, USA), MyD88, TRAF6, NF-κB (AF7524/AF8223/AF0246, Beyotime, China), p-NF-κB (TP56372, Abmart, China), GAPDH (TA309157, ZSGB-BIO, China) overnight at 4 °C and corresponding secondary antibody for 1h at room temperature. Finally, proteins were visualized using an enhanced chemical luminescent system (BLT GelView 6000 Pro).

### Cellular Metabolite Extraction and Metabolomics Profiling

The PA-stimulated cells were washed three times with precooled PBS. Then liquid nitrogen quenches the cells for one min. A total of 1 mL of precooled methanol-water (V: V=4:1, containing mixed internal standard, 4 μg/mL) was added to the sample and transferred to 2 mL centrifuge tubes in two separate aliquots. Then 200 μL of CHCl_3_ was added to the suspension. Samples were sonicated for 3 min, incubated on ice for 20 min, and left at -40 °C overnight to extract metabolites. The samples were centrifuged at 12000 × g for 10min at 4 °C. The supernatant was collected, dried under nitrogen, and finally re-extracted with 150 μL of mobile phase for LC-MS/MS analysis. Conditions for analysis of lipidomic by UPLC-Q mass spectrometry (Table S1). In order to investigate the stability of the instrument and the reproducibility of the samples, the quality control (QC) samples were prepared by pooling the same volume of each sample (Table S2).

The raw data were subjected to metabolomics processing software Progenesis QI v3.0 (Nonlinear Dynamics, Newcastle, UK) for baseline filtering, peak identification, integration, retention time correction, peak alignment and normalization. Compounds are identified based on multiple dimensions such as RT (retention time), exact mass number, secondary fragmentation and isotope distribution. The Human Metabolome Database (HMDB), Lipid Maps (v2.3), METLIN database and LuMet-Animal3.0 local database were used for identification and analysis.

## RESULTS

### Validation and LC-MS analysis of the analytical method

After the cellular sample was analyzed by 7-fold cross-validation, a principal component analysis diagram was generated. The fast contribution of the QC sample is shown in the diagram (Figure S1). The closely clustered QC samples indicate good stability and repeatability of the experiment. These results demonstrate the reliability and repeatability of the LC-MS analysis method shows that the control and model groups were completely separated (Figure S2).

### Metabolomic analysis of human macrophages subjected to PA treatment

The metabolites and lipids that made significant contributions to the difference between control and model groups were identified by S-plot of PCA and OPLS-DA analysis (Figure 2A and B). The volcano map clearly reflects 134 significantly up-regulated metabolites and 82 down-regulated metabolites in both groups (Figure 2C), indicating that the intervention of PA significantly altered the metabolites of macrophages. For cluster analysis, we selected 80 metabolites of interest. Compared with control groups, metabolites in the heatmap were significantly changed in the model groups (Figure 3A). Potential biomarkers identified in the database were used to analyze possible pathways through the KEGG database. KEGG enrichment analysis was performed on 80 differential metabolites, which were found to be up-regulated or down-regulated in 33 secondary pathways (Figure 3B). The top 20 pathways between the control and model groups are presented (Figure 3C and D). Among these, sphingolipid metabolism showed five differential metabolites (P<0.005), amino sugar and nucleotide sugar metabolism displayed six differential metabolites (P <0.005), and endocytosis demonstrated two differential metabolites (P <0.005).

**Figure 2.**
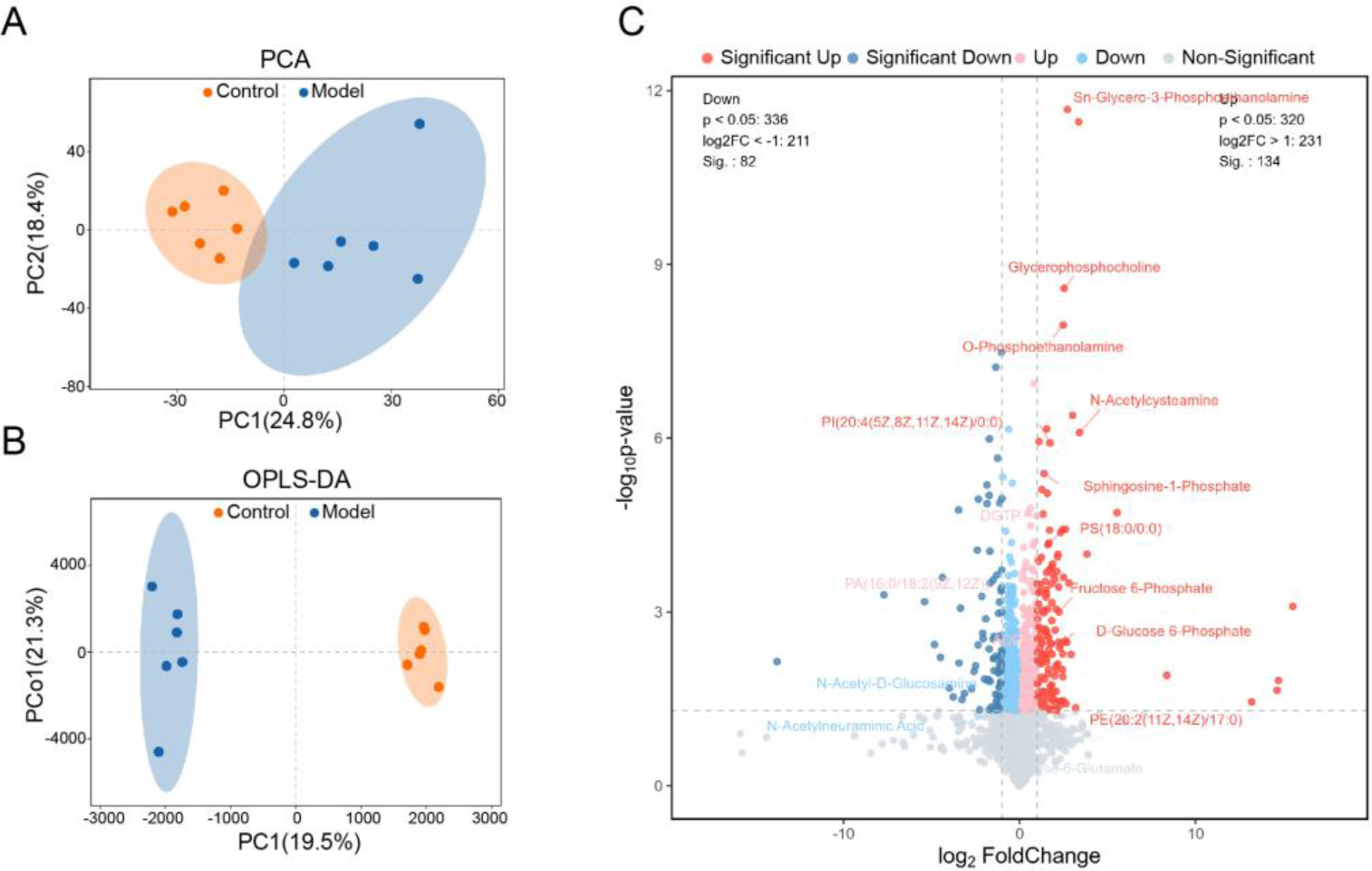
Multivariate analysis of cellular metabolomics data. (A) pca score plot of metabolomics in ESI+/ESI-ionization mode in the control and model groups. (B) OPLS-DA score plot of metabolomics in ESI+/ESI-ionization mode in the control and model groups. (C) Volcano plots of significantly upregulated and downregulated metabolites in the control and model groups.

**Figure 3.**
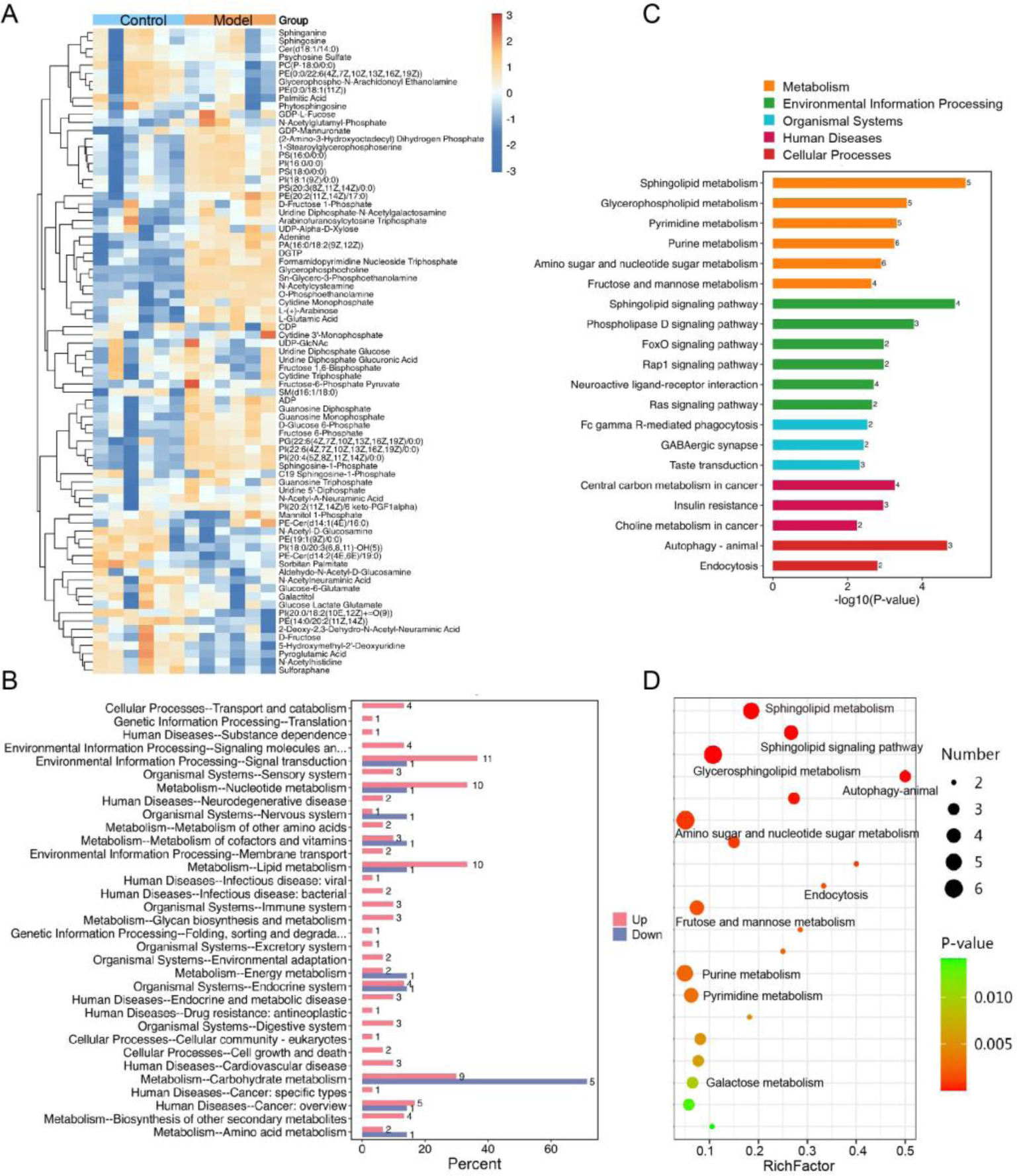
Metabolic effects of PA on metabolites in THP-1 macrophages. (A) Cluster heatmap of differentially metabolites in the control and model group; (B-D) KEGG enrichment analysis and essential pathways.

### The analysis of metabolic pathways

Amongst the cellular metabolites with the most significant changes associated with the TLR4 inflammatory pathway were mainly sphinganine-1P, sphingosine-1P and O-phosphoethanolamine (Figure 4A). Metabolites related to N-glycosylation substrates with alterations mainly include D-glucose 6-phosphate (Glc-6P), fructose 6-phosphate (Fruc-6P), GlcNAc, Neu5Ac, GDP-mannuronate, GDP-L-Fuc and UDP-GlcNAc/GalNAc (Figure 4B).

**Figure 4.**
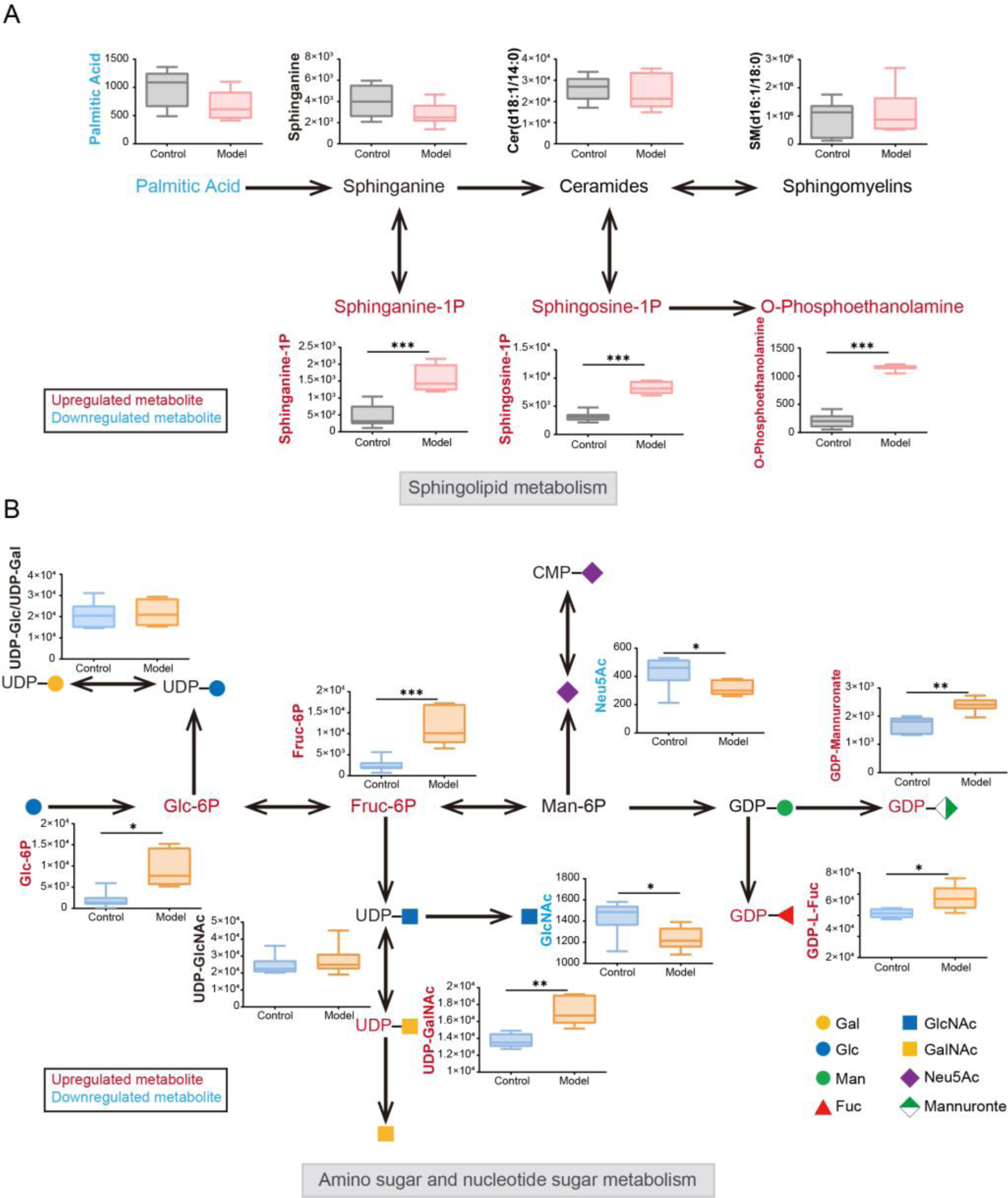
The relative abundance of potential biomarkers between the control and model groups. (A) Box plots of relative abundance for the seven sphingolipid metabolites; (B) Box plots of relative abundance for the nine amino sugar and nucleotide sugar metabolites. The upregulated metabolites are highlighted in red, while the downregulated metabolites are indicated in blue. Data are presented as mean ± SD (n=6) and were analyzed by ANOVA with Dunnett’s post-hoc analysis. *P<0.05; **P<0.01.

### Palmitic acid elicits TLR4-mediated inflammatory response in macrophages

We examined the effect of PA on TLR4/MyD88/TRAF6/NF-κB signaling pathway in macrophages at different time points. As expected, a time course study of TLR4/MyD88/TRAF6/NF-κB pathway protein levels in macrophages by Western blot, data analysis showed that treatment with PA induced MyD88 and TRAF6 expression and NF-κB phosphorylation, indicating canonical NF-κB activation (Figure 5A). After 12 hours of PA-incubated human macrophages, TLR4 expression was upregulated in (Figure 5B), and MyD88, TRAF6 and p-NF-κB were also upregulated, reaching the maximum at 24 hours (Figure 5C-E). The results of TNF-α and IL-1β assay in cell culture medium showed that the production of TNF-α and IL-1β in PA-stimulated macrophages gradually increased with time and then rapidly decreased after reaching the maximum at 36h (Figure 5F and G). Accordingly, in agreement with the data from the cell flow analysis of TLR4 endocytosis, the immune-fluorescent analysis suggested that TLR4 moved from the cell membrane to the interior of the cell and began internalization after 24 hours (Figure 6A). Antibodies recognizing TLR4 (both monomeric and dimeric TLR4) can be used to monitor TLR4 endocytosis in flow cytometry protocols^44,45^. We compared TLR4 endocytosis in palmitate-treated macrophages in various time points. TLR4 expression on the membrane decreased with increasing incubation time (Figure 6B-D), particularly after 24 hours (Figure 6E). Collectively, Palmitic acid-induced TLR4 endocytosis was detectable in macrophages during the experiment, these results provide further evidence that palmitate is induces trigger TLR4-mediated inflammatory response in macrophages.

**Figure 5.**
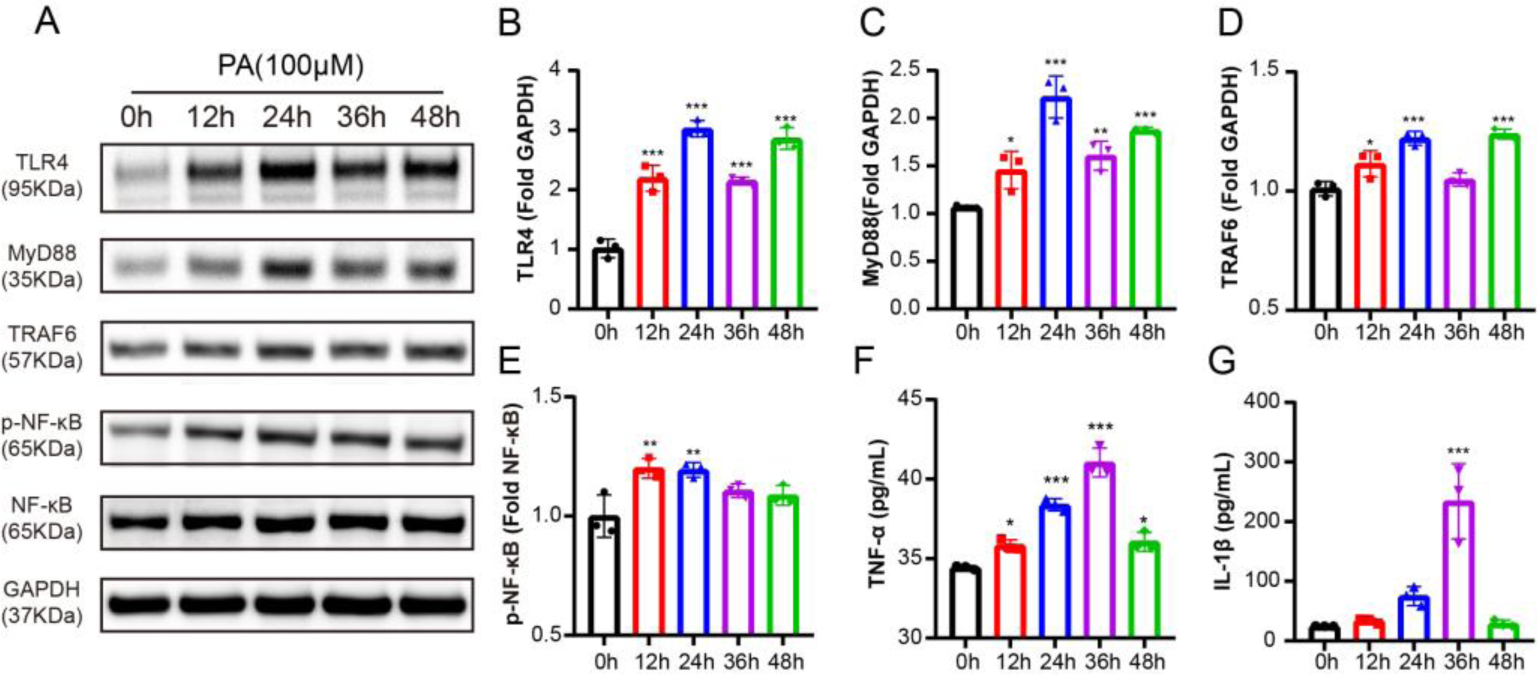
Changes of TLR4 signaling pathway on macrophages with PA stimulation time. THP-1 were pretreated with PMA(100 ng/ml) for 24 h to differentiate into THP-M. Then these cells were continuously stimulated with PA(100 μM). The levels of TLR4 signaling pathway in macrophages were determined using Western blotting (A). Western blots to detect TLR4 (B), MyD88 (C), TRAF6 (D), NF-κB and p-NF-κB (E) from macrophages treated with PA(100 μM) at 0 h, 12 h, 24 h, 36 h, 48 h. Meanwhile, the level of TNF-α (F) and IL-1β (G) in the cell supernatant were detected. Data are presented as mean±SD (n=3) and were analyzed by ANOVA with Dunnett’s post-hoc analysis. *P<0.05; **P<0.01; ***P<0.001.

**Figure 6.**
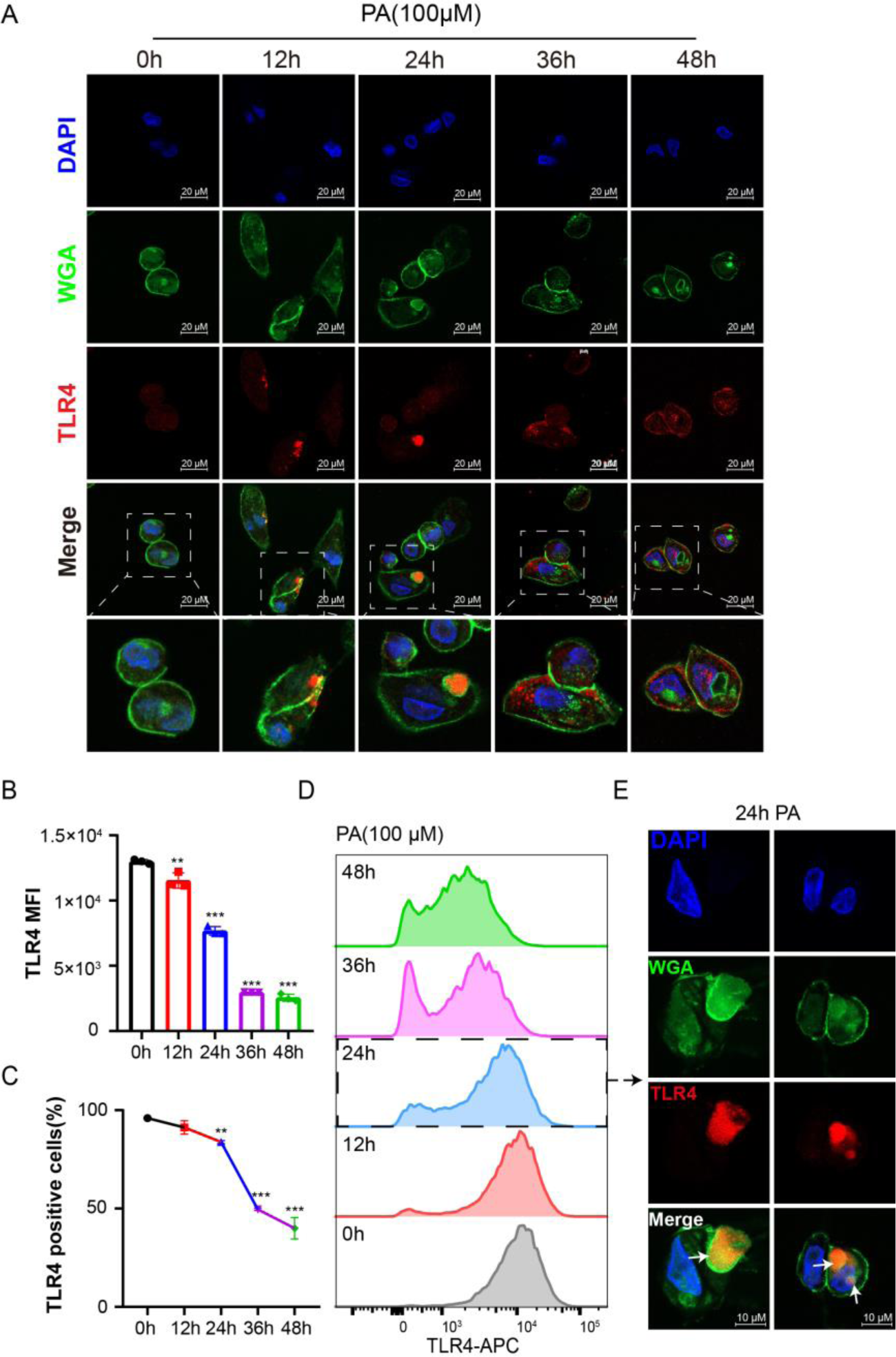
PA is able to induce TLR4 endocytosis. Macrophages were treated with PA (100 μM) for the indicated durations. Following stimulation, cells were fixed, stained with FITC-conjugated WGA (green), DAPI (blue ), anti-TLR4 antibody (red). Scale bars: 20 µm. (A) the level of TLR4 expression (B and D) and positive cell rate (C) were detected by flow cytometry. (E) Representative immunofluorescence images of TLR4 endocytosis in macrophages treated with PA for 24h. Scale bars: 10 µm. Data are presented as mean ± SD (n=3) and were analyzed by ANOVA with Dunnett’s post-hoc analysis. *P<0.05; **P<0.01; ***P<0.001.

### The impact of glycosylation on TLR4 activation in macrophages treated with PA

GalAz was introduced into the THP-1 cell culture system, followed by activation with PMA to induce macrophage differentiation. The Western blot results demonstrated that GalAz did not exert any discernible impact on TLR4 pathway in THP-1 macrophages (Figure S3). Subsequently, macrophages were incubated with PA and GalAZ at different time points and labeled with Fluor 488-alkyne (Figure 7A). Immunofluorescence analysis revealed an initial low expression of TLR4 in macrophages at 0 h, with predominant co-localization of TLR4 and GalAz observed on the cell membrane. After 12 hours, there was an increase in TLR4 expression in macrophages along with detection of TLR4-GalAz colocalization both in the cytoplasm and on the cell membrane. The expression of macrophage TLR4 continued to rise after 24 hours, accompanied by abundant cytoplasmic co-localization between TLR4 and GalAz. By 36 hours, dissociation between TLR4 and GalAz predominantly occurred, resulting in the transfer of TLR4 from the cell membrane to the cytoplasm (Figure 7B-D).

**Figure 7.**
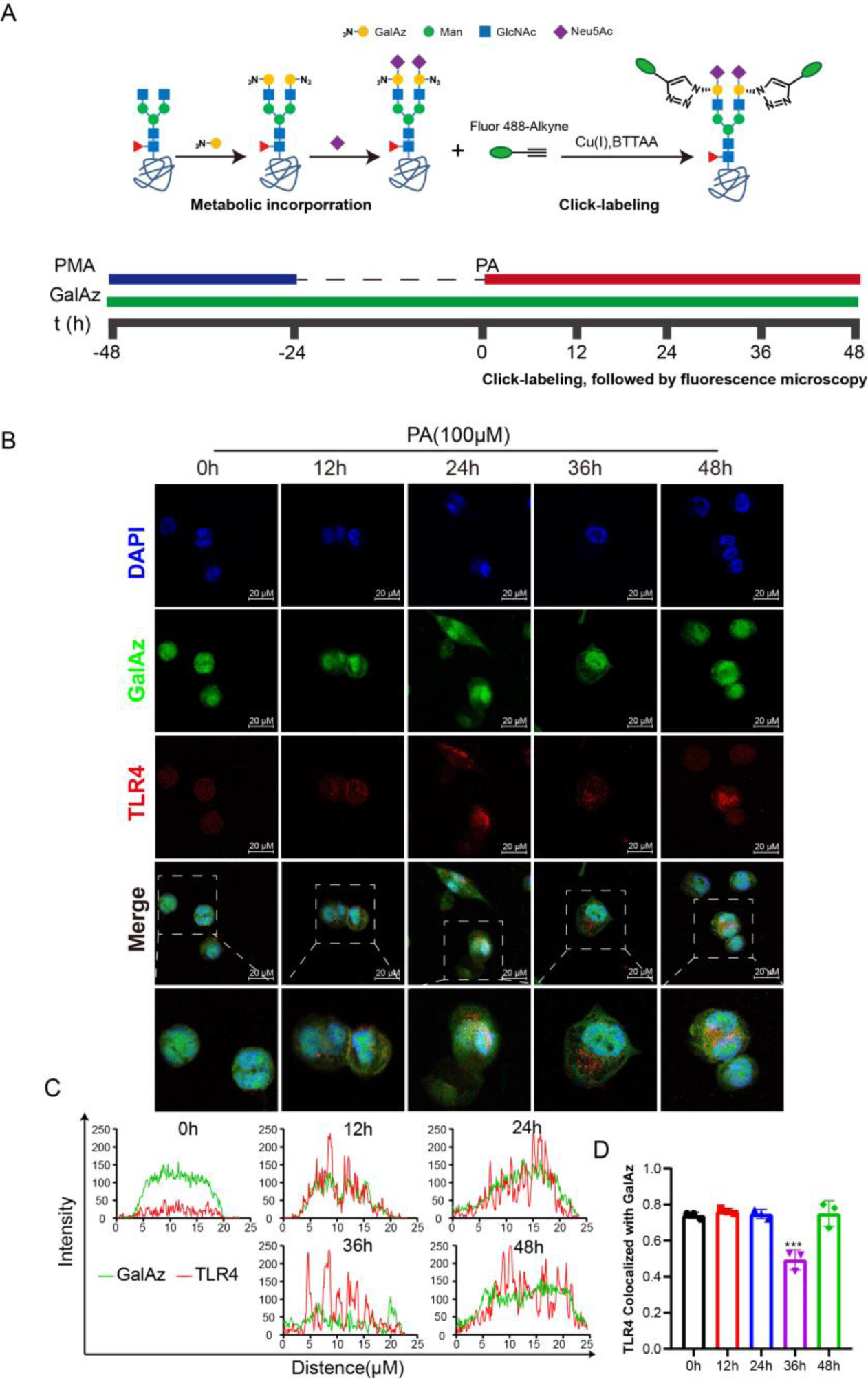
GalAz colocalizes with TLR4. (A) A schematic of the experimental procedures for metabolic labeling of GalAz glycans. After differentiation into macrophages, the cells were treated with PA (100 μM) for the indicated times; (B) Confocal fluorescence microscopy imaging of azide-labeled macrophages which stained with Fluor 488-alkyne(Green), DAPI (blue ), anti-TLR4 antibody (red). Scale bars: 20 μm. (C) Colocalization analysis of TLR4 and GalAz. (D) Pearson ’s Coefficient of TLR4 with GalAz is shown in bar graph format. Data are presented as mean ± SD (n=3) and were analyzed by ANOVA with Dunnett’s post-hoc analysis. *P<0.05; **P<0.01; ***P<0.001.

## DISCUSSION

Although substantial progress has been made in understanding the mechanism of fatty acids, especially PA, on the macrophage TLR4 inflammatory signaling pathway, studies on the N-glycosylation of TLR4 have also clarified the glycosylation process after the activation of TLR4. However, the establishment of macrophage metabolic biomarkers by high-throughput methods for the diagnosis and monitoring of macrophage TLR4-related inflammatory responses remains of great importance. To our knowledge, this is the first study to use metabolomics to identify potential biomarkers of PA-based macrophage inflammatory response.

A large number of studies have shown that free fatty acids^46–48^, especially saturated fatty acids ^49–51^, can activate TLR4 and its downstream signaling pathways, and trigger inflammatory responses in macrophages, of which PA is a typical representative^29^. Based on the effect of PA on human macrophages, we selected human macrophages cultured in PA for 24 h for metabolomic analysis. The metabolomic and lipidomic characteristics between Control and Model were significantly changed. Based on ROC analysis, sphingolipids metabolites (sphinganine-1P, sphingosine-1P and O-Phosphoethanolamine) and nucleotide sugar metabolites (Glc-6P, Fruc-6P, GlcNAc, Neu5Ac, GDP-mannuronate, GDP-L-Fuc and UDP-GlcNAc/GalNAc) provided effective diagnostic indicators. Therefore, it may be a new metabolic indicator of disease activity.

Sphingosine-1P, a sphingolipid metabolite downstream of ceramide, is important for inflammatory and immune responses. Sphingosine-1P has been implicated in many pathophysiological conditions and diseases, including inflammatory and autoimmune diseases^52^. In reactions catalyzed by the sphingomyelin kinases SPHK1 and SPHK2, sphingosine-1P is formed by the phosphorylation of sphingomyelin. SPHK1 can be activated by a variety of pro-inflammatory factors and promote the formation of Sphingosine-1P^53^. TLR4-mediated pro-inflammatory macrophage activation is characterized by increased glycolysis and alterations in mitochondrial metabolism, leading to the resynthesis of fatty acids that flow into sphingolipids, which have a regulatory role in the proinflammatory and pyrolytic phases of TLR4 activation in macrophages^54^. Lipopolysaccharide (LPS) and palmitate can activate the TLR4-mediated inflammatory signaling pathway and lead to a synergistic increase in the levels of Sphingosine-1P^55,56^. And sphingosine-1P has been shown to be an activator of the key inflammatory transcription factor NF-κB^57^. In our study, sphingosine-1P was upregulated in the PA cohort. The upregulation of sphingosine-1P may be part of the explanation for PA incubation of macrophage inflammatory response. In support of the existing findings, using several independent experimental approaches, we have provided evidence that PA could upregulate the expression of TLR4 in human macrophages and activate the TLR4 signaling pathway.

Here, employing multiple independent experimental approaches, e present compelling evidence demonstrating that PA activates TLR4 and induces inflammation in macrophages via the MyD88/TRAF6/NF-κB signaling pathway. From our results, we can see that when macrophages were incubated with PA, the expression of TLR4 protein tends to increase, whereas the expression of MyD88, TRAF6, NF-κB, and p-NF-κB proteins, which correspond to the TLR4 inflammation pathway, did not change obviously. 24 h later, TLR4, MyD88, TRAF6, NF-κB and p-NF-κB protein expression were significantly upregulated. This indicates that human macrophages incubated with PA showed a progressive increase in TLR4 expression before 12 h, and the transduction of TLR4 inflammatory signals became active after 12 h. Although current studies are inconsistent as to whether the effect of PA on TLR4 is direct or indirect, it is well established that PA can activate the TLR4/MyD88/TRAF6/NF-κB signaling pathway and promote the release of inflammatory cytokines in macrophages^27^.

The formation of the TLR4/MD-2 heterotetramer is the most recent event to initiate the TLR4 pathway^45^. However, the activation induced dimerization of TLR4 is internalized into endosome through endocytosis, which is the key event for TLR4 pathway activation^44^. Our results suggest that TLR4 is activated on the membrane of human macrophages incubated with PA for 12 hours. Significant internalization of TLR4 was observed after 24 hours. The expression of TLR4 in the cell membrane decreased after 24 hours and was abundant in the cell interior. Thus, activation by dimerization of the active TLR4/MD-2 complex is followed by subsequent endocytosis leading to its translocation from the plasma membrane to endosomes. This further suggests that PA incubation of human macrophages activates TLR4 on the cell membrane, leading to an inflammatory response in the cells.

According to previous studies, TLR4 is a transmembrane protein that is initially localized in the ER. The surface expression of TLR4 is regulated by chaperone glycoprotein 96 (gp96) and protein associated with Toll-like receptor 4 (PRAT4A) ^58,59^. Gp96 is associated with immature TLR4 in the ER and induces the association of MD-2 with TLR4 to form TLR4/MD-2 complex^60^. In turn, PRAT4A is associated with the hypoglycosylated immature TLR4 to regulate TLR4 maturation. After PRAT4A interacts with TLR4, TLR4 becomes glycosylated and matures in the ER and Golgi^32,61^. The receptor substrate for N-glycosylation is the asparagine residue present in the consistent sequence N-X-S/T^62^. N-glycans are composed of Man, GlcNAc, Gal, GalNAc, Fuc and Neu5Ac^63^. During N-glycosylation, Fuc binds to the first GlcNAc that binds to asparagine, Gal and GlcNAc section at the end of the union, then may be Neu5Ac combined with the Gal^64^. Glc is the primary carbon source for humans and most organisms. From this source, the other monosaccharides required for glycan biosynthesis can be synthesized. The presence of glucose in the cell culture medium has a detrimental effect on the use of fluorescently labeled glucose for the monitoring of cell UDP-Glc dynamics. It is also interesting to note that inhibition of Neu1, either by gene deletion or administration of sialidase inhibitors, confers strong protection against endotoxemia^65^. Fluorescently labeled sialic acid tracing was therefore also excluded. Galactose acts as an N-glycosylated terminal linked sialic acid block, and its nucleotide sugar substrate UDP-Gal can be synthesized from galactose. We observed persistent co-localization of Gal and TLR4 in the membrane of PA-stimulated macrophages for 12 to 24 hours, which gradually shifted towards the cytoplasm. Based on our immunoblotting results, we propose that this phenomenon is attributed to nascent TLR4 glycosylation^66^. The eventual loss of Gal and TLR4 co-localization in the cytoplasm after 36 hours may be associated with lysosomal degradation following endocytosis. This inference is consistent with another study showing that N-glycosylation of TLR4/MD-2 is essential for its membrane expression and dimerization^33^.

This study found the levels of UDP-GlcNAc/UDP-GalNAc and GDP-L-Fuc were strongly increased in PA treated human macrophages. Fruc-6P, a glucose metabolite for the synthesis of UDP-GlcNAc/UDP-GalNAc and GDP-L-Fuc^66^, was also significantly increased. Nucleotide sugars are substrates for glycosylation (i.e. UDP-GlcNAc, GDP-L-Fuc, CMP-Neu5Ac, etc.) and are typically derived from glucose metabolism. N-glycosylation required GlcNAc, Gal and Fuc were produced by the cytoplasm of UDP-GlcNAc, UDP-Gal and GDP-L-Fuc. And Neu5Ac was converted from CMP-Neu5Ac synthesized by the nucleus^67^. The shift of these metabolites combined with the change trend of TLR4 expression at the corresponding time points suggested that the synthesis of glycosylation substrate nucleotide sugar was enhanced due to the requirement of N-glycosylation in the expression and activation of TLR4. The increase of Neu5Ac may be related to the synthesis of CMP-Neu5Ac and the endocytosis of TLR4 dimer after activation.

Macrophages play a central role in the host response to infection and tissue injury, and there is increasing evidence that these cells are dysfunctional in patients with obesity. Here, we describe the induction of TLR4 and it signaling by PA in human macrophages and observe that N-glycosylation is an important marker of TLR4 activation. At present, although the research on the biomarkers of TLR4 activation has made some progress, they mainly focus on the effect of LPS on macrophage TLR4, and the metabolites are limited to the de novo synthesis of fatty acids and the formation of complex lipids that construct proteins^54,68^. The metabolites related to N-glycosylation of TLR4 due to fatty acid activation are lacking. Our study started from the N-glycosylation of TLR4 and found that the metabolic molecule UDP-GlcNAc/UDP-GalNAc and GDP-L-Fuc could be a metabolic biomarker for TLR4 activation by fatty acids. Metabolic biomarkers UDP-GlcNAc/UDP-GalNAc and GDP-L-Fuc in obesity and related diseases diagnosis significance needs further research.

## Abbreviations

FFA: Free fatty acids
PA: Palmitic acid
TLR4: Toll-like receptor 4
ER: Endoplasmic Reticulum
NF-κB: Nuclear Factor-κB
MD-2: Myeloid Differentiation protein-2
Gal: Galactose
GalNAc: N-acetylgalactosamine
Glc: Glucose
GlcNAc: N-acetylglucosamine
Fuc: Fucose
Fruc: Fructose
Neu5Ac: N-Acetylneuraminic acid
UDP: Uridine diphosphate
GDP: Guanosine Diphosphate
CMP: Cytidine Monophosphate
GalAz: 2-azide-2-deoxy-D-galactose
S1P: Sphingosine 1Phosphate
SPHK1/2: Sphingomyelin kinase1/2
Gp96: Glycoprotein 96
MGAT1: alpha-1,3-mannosyl-glycoprotein 2-beta-N-acetylglucosaminyl transferase
LPS: Lipopolysaccharide
TNF-α: Tumor necrosis factor alpha
TRAF6: TNF receptor associated factor 6
MyD88: Myeloid differentiation primary response protein 88
NF-κB: Nuclear Factor-κB

## Ethics approval and consent to participate

All procedures performed conforming to the National Institutes of Health guide for the care and use of laboratory animals (NIH Publications No. 8023, revised 1978) and the Animal Management Rules of the Ministry of Health of the People’s Republic of China (No. 55, 2001), and the protocol was previously approved by the Institutional Ethics Committee for Use of Animals at Third Military Medical University (Chongqing, China).

## Availability of data and materials

Not applicable.

## Consent for publication

Not applicable.

